# Replication-related control over cell division in *Escherichia coli* is growth-rate dependent

**DOI:** 10.1101/2021.02.18.431686

**Authors:** Sriram Tiruvadi-Krishnan, Jaana Männik, Prathitha Kar, Jie Lin, Ariel Amir, Jaan Männik

## Abstract

How replication and division processes are coordinated in the cell cycle is a fundamental yet poorly understood question in cell biology. In *Escherichia coli* different data sets and models have supported a range of conclusions from one extreme where these two processes are tightly linked to another extreme where these processes are completely independent of each other. Using high throughput optical microscopy and cell cycle modeling, we show that in slow growth conditions replication and division processes are strongly correlated, indicating a significant coupling between replication and division. This coupling weakens as the growth rate of cells increases. Our data suggest that the underlying control mechanism in slow growth conditions is related to unreplicated chromosome blocking the onset of constriction at the midcell. We show that the nucleoid occlusion protein SlmA does not play a role in this process and neither do other known factors involved in positioning bacterial Z-ring relative to the chromosome. Altogether this work reconciles different ideas from the past and brings out a more nuanced role of replication in controlling the division process in a growth-rate dependent manner.

## Introduction

The studies addressing coordination between DNA replication and cell division cycles in *Escherichia coli* date back more than half a century and are still being strongly influenced by the classic Cooper-Helmstetter (CH) model [1]. The latter postulates that cell division completes a period of constant duration, referred to as D-period, after the termination of replication. The model also proposed that the replication period C is growth rate independent in a range of faster growth rates. A constant D-period, which was found to be independent of the growth rate in faster growth rates, would imply that cell division is tightly coupled to replication termination. These predictions have been revisited more recently using single-cell measurements. A good match between the CH model and data was found but only under a further assumption that the C+D period depends on the growth rate in a specific way [2].

As a new element for cell cycle control going beyond the CH model, the adder concept has been introduced [3–7]. In the adder model cells add a constant volume increment during the cell cycle irrespective of their size at birth. In some of these models, the increment is assumed to be added between two consecutive replication events, and cell division is still thought to be tightly coupled to replication [3, 6, 7]. In others, the increment is added from cell birth to division [4, 5, 8]. In these latter models, replication does not play any role in the division process. The latter conclusion has also been drawn by some experimental works which have not relied on cell cycle modeling [9, 10]. As the middle ground of these opposing views, Micali *et al*. have proposed a model postulating that division is controlled concurrently by replication and division-related processes; whichever of these processes completes the latest will trigger cell division [11, 12]. In this concurrent-processes model replication and division related processes are competing with each other in triggering cell division in all growth conditions.

Cell division occurs when two daughter cells separate from each other. This event can be determined from single-cell time-lapse measurements. The existing models predict the timing and cell sizes at this event. However, it is well-known that the separation of daughter cells results from a long sequence of biochemical processes that only culminate with the separation of two daughter cells [13, 14]. The question arises on what initiates this process sequence and how this initiation is linked to the replication cycle of the chromosome. Previous research has identified that cell division in *E. coli* progresses via two distinct stages [15]. The first of these is the formation of the Z-ring at the cell center. This comprises the assembly of FtsZ protofilaments in the mid-cell region [16]. In *E. coli* the protofilaments are linked to the cell membrane by FtsA and ZipA linkers and likely bundled together with ZapA and several other cross-linking proteins [14, 16]. At multi-forked fast growth conditions, the Z-ring forms at cell birth but in slower growth conditions there is a delay between cell birth and Z-ring formation, which is at least in part controlled by the availability of FtsZ [17, 18]. The Z-ring subsequently recruits about 30 different proteins that are involved in septal cell wall synthesis and partitioning of DNA between daughter compartments [19, 20]. The recruitment of these mostly regulatory proteins proceeds in specific order culminating with the recruitment of FtsN to the divisome complex [14, 16, 19, 21–23]. It is hypothesized that FtsN relieves inhibition or activates core septal peptidoglycan synthesis complex consisting of transpeptidase (FtsW) and transglycosylase (FtsI/PBP IIIA) units [24]. There is a significant delay between the formation of the Z-ring and the onset of constriction [15]. The latter was found to be simultaneous with the recruitment of FtsN to the divisome [22]. It is currently unclear why Z-rings form much earlier than constriction is initiated. Furthermore, the experimental studies related to molecular aspects of cell division have not addressed the question of how the recruitment of divisome components is regulated by the replication cycle of the chromosome.

Here, we study how the initiation of constriction is controlled by the replication cycle. We use quantitative high-throughput fluorescent microscopy and a new functional endogenous FtsN construct. The latter allows us to accurately determine the timing for constriction formation. Our data and cell cycle modeling are consistent with an idea that replication is a rate-limiting factor for constriction in slow growth conditions. However, the limiting role of replication weakens at faster growth rates. Our results furthermore suggest that the onset of constriction is limited by the unreplicated chromosome at the midcell. This limitation is not related to the nucleoid occlusion factor SlmA, the Ter linkage proteins (ZapA, ZapB, and MatP), and FtsK, a DNA translocase.

## Results

### Constriction formation follows replication termination in different growth conditions

To understand the link between replication and division cycles we constructed *E. coli* strains where fluorescent fusion proteins labeled both the replisome and the divisome (for details see Materials and Methods, SI Table S1). We used the N-terminal fusion of mCherry to DnaN (beta clamp) [25] or C-terminal fusion of Ypet to ssb (single-strand binding protein) to label the replisome [26]. For the divisome label, we chose FtsN because it is the latest known component to assemble to the divisome and its recruitment has been reported to coincide with the onset of constriction [14, 16, 19, 21–23]. While in previous fluorescent constructs of FtsN the labeled protein was expressed from extra copy plasmids [19, 27, 28], in our construct it was expressed from the native locus. We grew these strains in steady-state conditions in mother machine devices [29, 30]. The doubling times and lengths of these cells were indistinguishable from the WT ones (strain BW27783) when grown in a glycerol medium (Table S2). Note that all measurements were performed at 28 °C where the growth rate is expected to be about 2 times slower than at 37°C [31]. Using the fluorescently labeled strain we followed the timing of replication termination (*Trt*), onset of FtsN accumulation at mid-cell (*Tn*) and onset of constriction (*Tc*) in time-lapse images (Fig. 1A-B). Here all the times are given relative to cell birth. Additionally, we also determined the timing of replication initiation (*Tri*) and the C-period (*C* = *Trt* – *Tri*). We determined *Tri, Trt* and *Tn* from the analysis of fluorescent images and *Tc* from the phase images (for details see Methods). We found *Tc* to be delayed relative to *Tn* on average by about 12 mins (SI Fig. S1). We assign the delay to less sensitive determination of constriction formation from phase images. We therefore use *Tn* instead of *Tc* for the timing of the constriction formation in the Figures in the main text while the data on *Tc* can be found in SI Figures.

**Figure 1.**
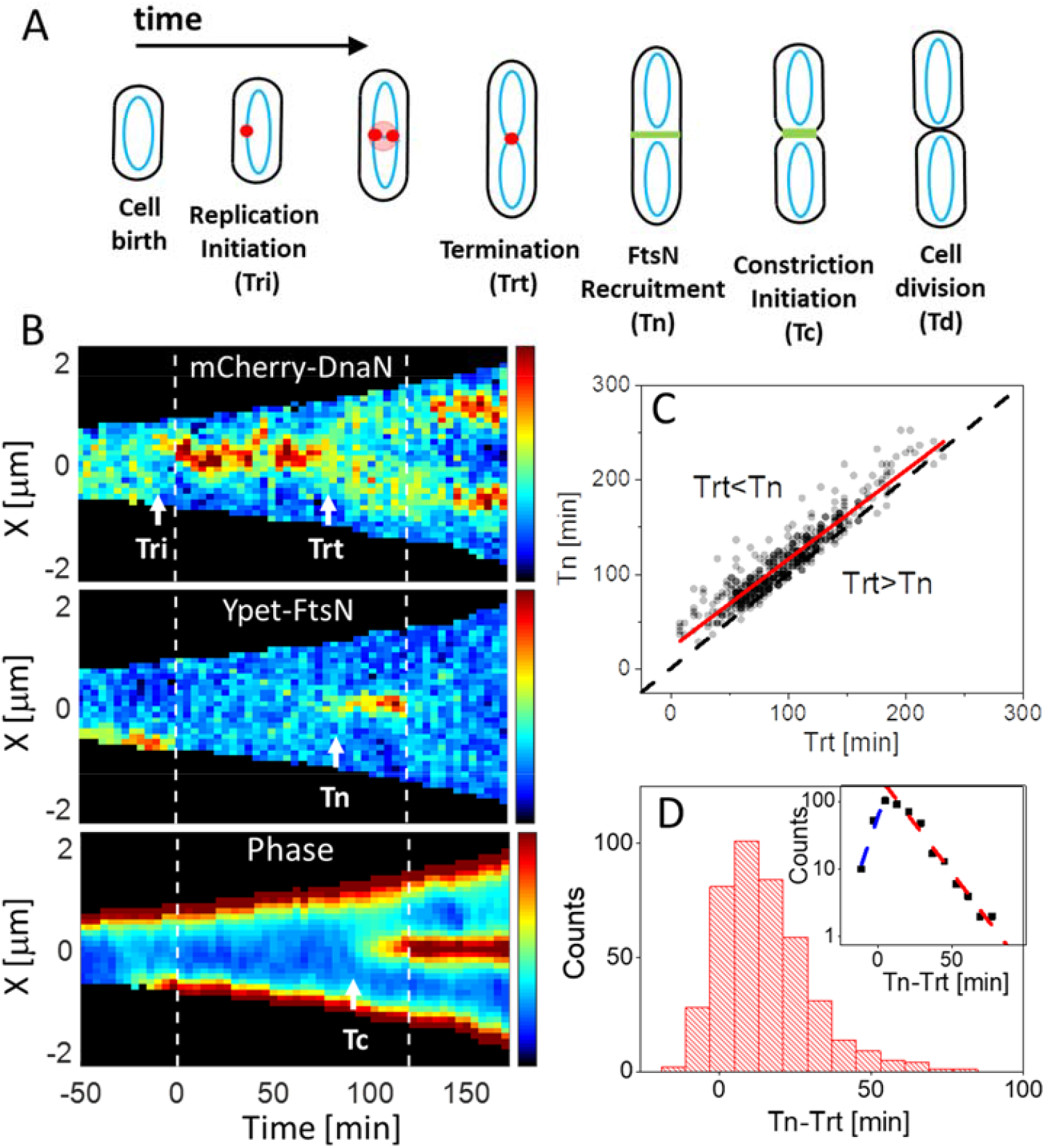
Timing of constriction formation and recruitment of FtsN relative to termination of replication in slow growth conditions. (A) Schematics for the main cell cycle events and timings that are determined from time-lapse measurements. (B) Kymographs of fluorescent and phase signals for a representative cell grown in M9 glycerol medium. Dashed vertical lines indicate cell division events. Red corresponds to high and blue to low-intensity values. Event timings are indicated by arrows. (C) Termination of replication (*Trt*) vs initiation of constriction (*Tn*) for a population of cells (N=420). *Tn* is determined based on the accumulation of Ypet-FtsN signal at mid-cell. The solid red line is a linear fit with (*Tn* = 0.94*Trt* + 22 mins). The dashed black line corresponds to *Trt* = *Tn*. (D) Distribution of delay times between constriction formation and termination of replication for these cells. Inset shows the same data in a semi-logarithmic plot. The dashed lines are fits to exponential decay. The time constant for the fit at negative times is 7 min and for positive times 15 mins.

We first investigated the correlation between termination and onset of constriction times in slow growth conditions in the M9 glycerol medium (Fig. 1C). The *Tn* and *Trt* times were correlated (with a Pearson correlation coefficient *R* = 0.94) as were also *Tc* and *Trt* times (*R* = 0.92; SI Fig. S1). The comparable timings and correlations between *Tc* and *Trt* time were also present in a different strain which carried ssb-Ypet label for replisome and no divisome label (Fig. S2) indicating that Ypet fusion to FtsN and mCherry fusion to DnaN did not have significant effects on division and replication processes.

In 7% of cells, we found the onset of constriction as measured by Ypet-FtsN (i.e., *Tn*) occurred before the termination (*Trt*) (Fig. 1D). When we determined the onset of constriction from the phase images (*Tc*), in only 1 out of 420 cells termination occurred earlier than the onset of constriction (SI Fig. S1) but as argued earlier the latter estimate is likely less accurate. For 7% of cells, in which the initiation of constriction preceded the termination, the distribution of times *Tn* – *Trt* was approximately exponential with a characteristic time of 7 min (inset of Fig. 1D). The latter time is close to the characteristic time that DnaN remains attached to the replication terminus region after completion of replication (3 mins at 37 °C, potentially translating to about 6 mins in our conditions) [25]. Altogether, the fraction of cells in which the termination of replication occurs after the *actual* onset of constriction is much smaller than 7%, if not zero. Interestingly, the distribution of *Tn* – *Trt* was also approximately exponential for positive values with a characteristic time of 15 mins (Fig. 1D, inset). The latter suggests the possibility of a single rate-limiting reaction associated with the process of triggering constriction formation that follows the termination, as we elaborate on later.

We next investigated how the above conclusions applied at different growth rates. We repeated these measurements in eight additional different growth media (SI Table S3); in three of these, the growth rates were slower while in the other five the rates were higher than in the measurement discussed above (SI Table S2). In all of these nine growth conditions, the average delay time ⟨*Tn* – *Trt*⟩ was positive showing that constriction formation follows on average the termination and possibly in all divisions (Fig. 2A, inset). ⟨*Tn* – *Trt*⟩ showed variation between 20 min to 40 mins in different growth rates except for the slowest growth rate in acetate medium where ⟨*Tn* – *Trt*⟩≈ 65 *min*. Unlike the almost growth rate-independent behavior of ⟨*Tn* – *Trt*⟩, the normalized delay times, ⟨(*Tn* – *Trt*)/*Td*⟩, showed two distinct growth-rate dependent regimes (Fig. 2A). Below about *Td* ≈130 mins the normalized times decreased as the doubling time increased but above it, the values plateaued reaching about 12% of the cell cycle. A similar cross-over from one regime to another was also seen in Pearson correlation coefficients, *R*(*Tn, Trt*) (Fig. 2B). For *Td* > 130 mins the termination of replication and the onset of constriction were highly correlated (*R*(*Tn, Trt*) > 0.85) and independent of *Td*, while for *Td* < 130 mins these correlations decreased approximately linearly with the decreasing *Td*. A similar cross-over behavior could be also seen in plots when the timing of constriction (*Tc*) was determined from phase images (SI Fig. S3).

**Figure 2.**
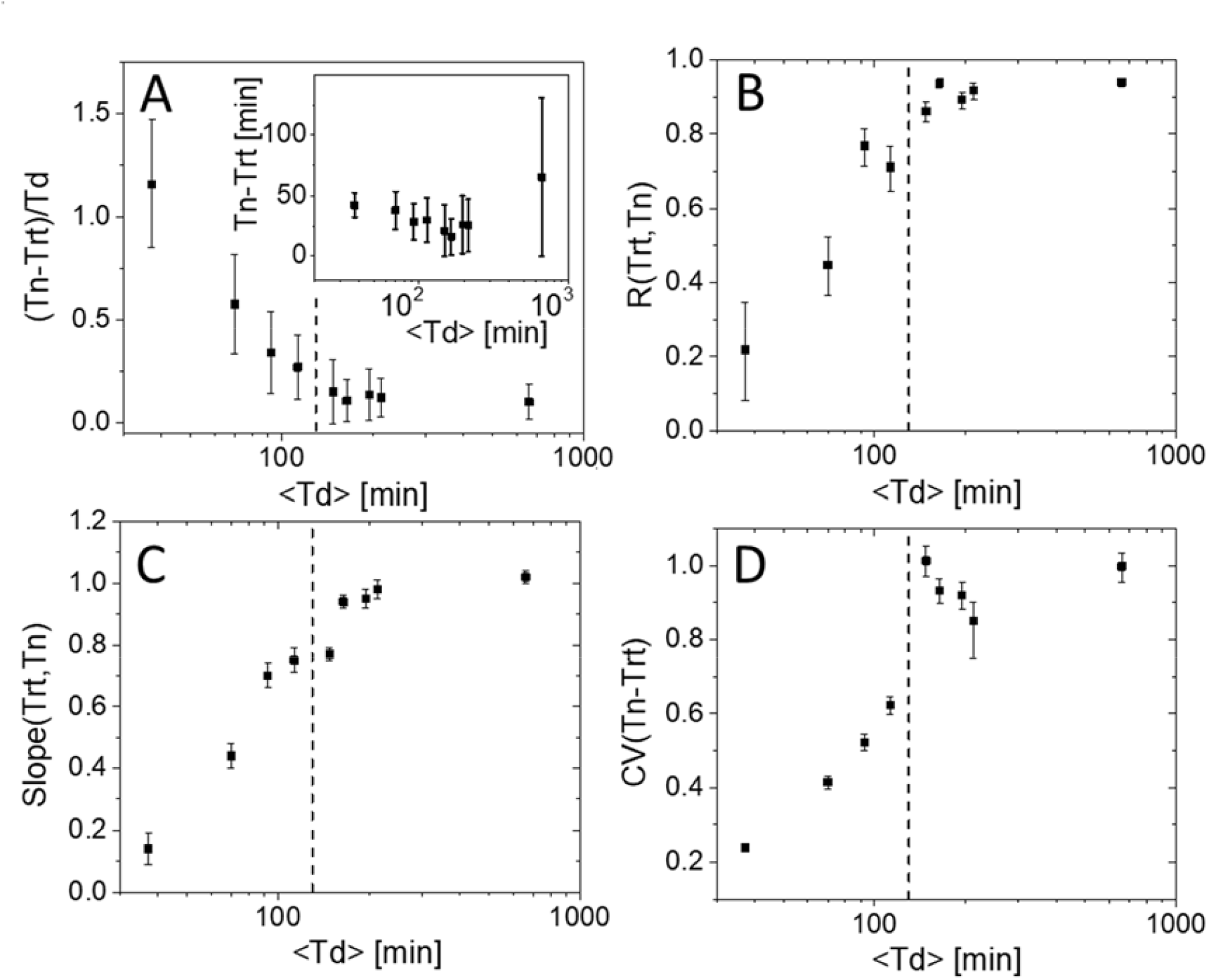
Comparison of timings of constriction initiation and termination of replication in 9 different growth media. From the longest to shortest doubling times the carbon sources used in the media are acetate, alanine, mannose, glycerol, glycerol + trace elements (TrEl), glucose, glycerol+Cas, glucose+Cas, and EZ-Rich defined medium with glucose (for details see Table S3). (A) The average normalized delay time between initiation of constriction and termination of replication as a function of the average doubling time, ⟨*Td*⟩. Inset shows the unnormalized delay time. Error bars in both plots show the std of these quantities within the cell population. (B) Pearson correlation coefficient between *Trt* and *Tn*. (C) The slope of *Trt* vs *Tn* plot. (D) Coefficient of variation for *Trt* – *Tn* distribution. The dashed vertical lines in all plots correspond to ⟨*Td*⟩ = 130 min. Error bars in (C)-(F) show 95% confidence intervals. For calculation of these intervals see Methods.

The times of termination and constriction initiation were not only correlated but furthermore followed a timer-like relationship, *Tn* = *Trt* + *constant*, at slower growth rates. This was evident in plots of *Trt* vs *Tn* where linear regression gave a slope of ≈1 for longer doubling times (Fig. 2C). The corresponding intercept of the fits was almost independent of the doubling time (SI Fig. S4A). We also found that the distribution of delay times *Tn* – *Trt* in a given growth condition was approximately exponential at slower growth rates, as it was for the growth condition described above (SI Fig. S4B). This was also reflected in the coefficient of variation (CV) of these distributions, which was approximately one at longer doubling times (Fig. 2D). The CV values also showed a cross-over at *Td* ≈130 mins. In shorter doubling times, the CV values decreased and the mode of the *Tn* – *Trt* distributions shifted to positive values (SI Fig. S4B).

Altogether, the exponential distribution of delay times and the timer behavior in a range of slow growth conditions suggest a constant rate process linking onset of constriction to replication. The process may result from a single first-order reaction with some rate-limiting component. Irrespective of the details of this process, for *Td* > 130 mins our data is consistent with the idea that some replication related process controls the initiation of constriction but as the doubling times shorten this process becomes less and less rate-limiting.

### Model supports checkpoint for constriction to be close to termination in slow growth conditions

The presented data in slow growth conditions suggest that there is a replication-dependent checkpoint for the onset of constriction. However, it may occur before the termination of replication. To narrow down the possible time range for this checkpoint we constructed an analytical model. The model allows us to calculate the *Tn* – *Trt* distributions and various statistics related to the processes as a function of the timing of the checkpoint, *Tx* (Fig. 3A). In the following discussion, we define the normalized time difference *x* = (*Trt′* – *Tx*)/⟨*C*⟩, namely the time delay between the true termination event, *Trt* (as opposed to the *measured* termination event *Trt*) and the checkpoint *Tx*. In the above expression ⟨*C*⟩ is the average C-period in a given growth condition. *x* ranges from 0 to 1, with *x* = 0 corresponding to a checkpoint at the termination and *x* = 1 to one at the initiation of replication. We furthermore assume that when the replication fork reaches a relative distance *x* from the replication terminus, initiation of constriction occurs with a constant rate *r* (i.e., consistent with first-order reaction kinetics). The assumption of a single rate constant is based on the approximately exponential distribution for *Tn* – *Trt* (for positive values) (Fig. 1D, SI Fig. S4B) as well as the CV of this distribution being approximately equal to 1 (Fig. 2E) in slow growth conditions. The model also accounts for the difference between the measured value of *Trt* and the actual one (*Trt*′) due to the finite time DnaN remains DNA bound after replication completes. The attachment time of DnaN has been found to be exponentially distributed [25]. As discussed previously, a mean time of ⟨*Ta*⟩ = 3 to 6 *mins* can be expected in our growth conditions. Under these assumptions, we find that the CV of the *Tn* – *Trt* distribution is given by:

**Figure 3:**
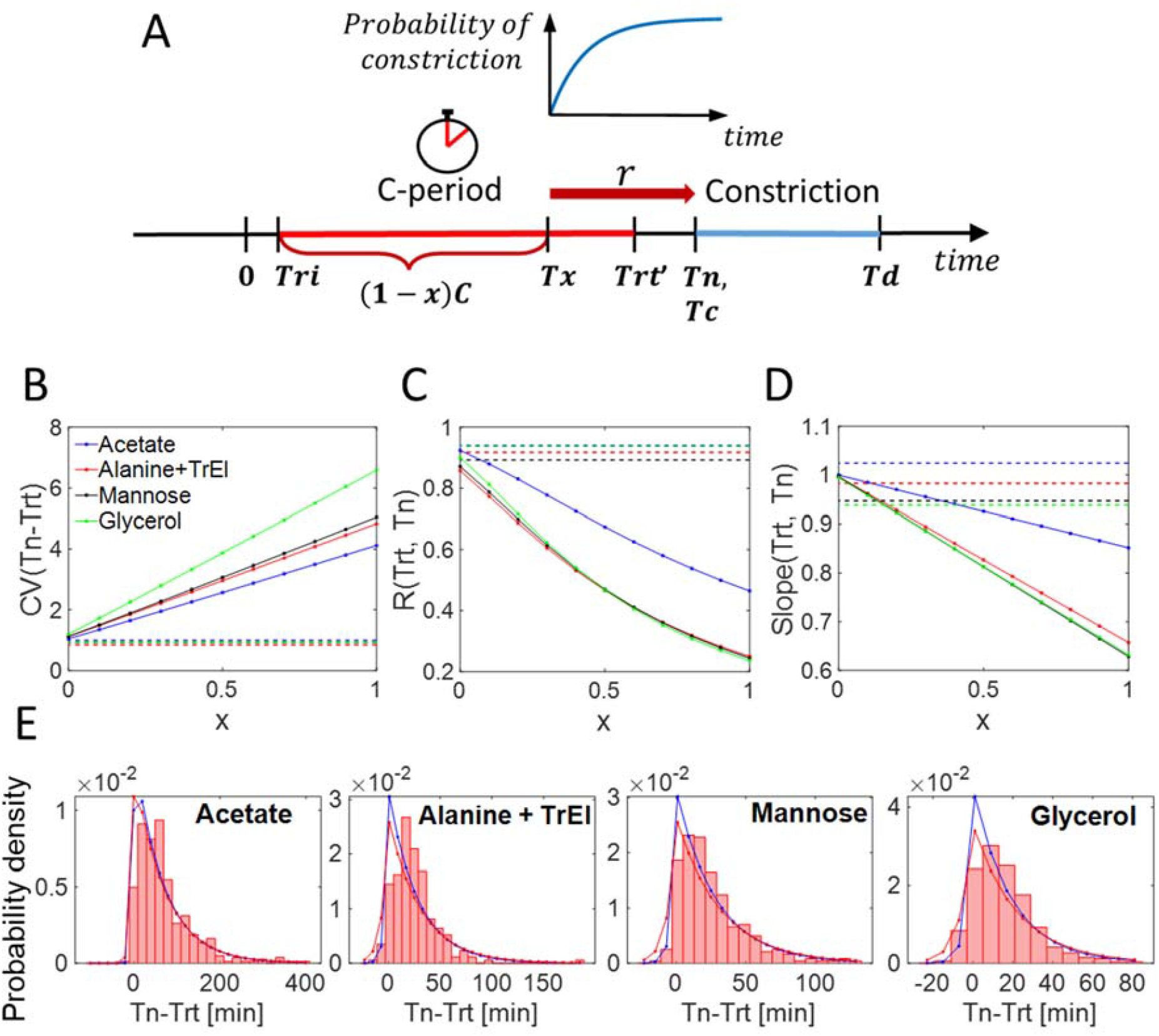
Predictions of model coupling the replication cycle to the onset of constriction. (A) Schematics representing the model. *Tx* is the timing for the checkpoint that triggers constriction formation. *x* is the normalized time of this checkpoint from termination. *Trt*′ is the actual time of termination, which differs from the measured time *Trt* by the detachment time of mCherry-DnaN from the chromosome (see Methods for details). (B) Coefficient of variation of the *Tn* – *Trt* distribution, (C) Pearson correlation coefficient between *Trt* and *Tn*, and (D) the slope of the linear regression line for *Tn* vs *Trt* all plotted as a function of *x*. In panels (B)-(D) the solid lines show predictions of the model and the dashed horizontal lines the experimental values. Note that only the four slowest growth conditions are considered in these comparisons and ⟨*Ta*⟩ = 3 *min*. (E) Distribution of *Tn* – *Trt* for slow-growth conditions obtained from experiments and from theory for two different values of ⟨*Ta*⟩. The theoretical distributions are given by eq. 20 in Methods and they correspond to *x* = 0. Blue lines correspond to ⟨*Ta*⟩ = 3 *min*, and red lines to ⟨*Ta*⟩ = 6 *min*. All other parameters in eq. 20 are determined from the data.

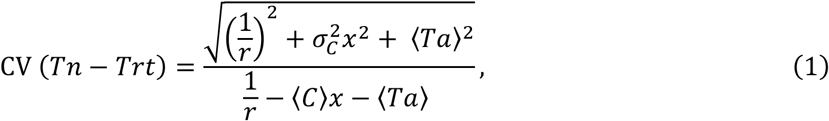

(see Model in Methods for the derivation). Here, *σ*_C_ is the standard deviation for the distribution of C-periods within the cell population. All the quantities except *x* in eq. 1 (namely ⟨*C*⟩, *σ_C_, r*, ⟨*Ta*⟩) are determined from experiments (see Model in Methods). We compared the predictions of the above formula to the measured CV values in the four slowest growth conditions. Eq. 1 predicts that *CV*(*Tn* – *Trt*) is close to 1 near *x* = 0 and rises rapidly with increasing *x* for all growth conditions considered (Fig. 3B). The experimentally measured CV values are thus consistent with the model only when *x* is close to zero, that is when the constriction is initiated shortly before or at termination.

The Pearson correlation coefficients between *Tn* and *Trt* (Fig. 3C) and the slopes of *Tn* vs *Trt* linear fits (Fig. 3D), both of which can be derived analytically within the model, also show an agreement with the experimental data only when *x* is close to zero. In some growth conditions, however, the best agreement between model and data for the slope of *Tn* vs *Trt* occurs for larger values of *x*. The most outlying point in Fig. 3D corresponds to mannose where the data and the model agree at *x* = 0.2. Nonetheless, most data appear to be consistent with the checkpoint at the termination. Indeed, taking *x* = 0, we could explain satisfactorily the entire distribution of *Tn* – *Trt* for all slow growth conditions (Fig. 3E).

To further test the model, we also compared correlations and statistics between experimentally determined replication initiation, *Tri* and constriction initiation, *Tn*, timings (SI Fig S5 A-D, Table S4). We found that the model agrees with the experimental values of CV(*Tn* – *Tri*) and *R*(*Tri, Tn*) in all slow growth conditions for *x* < 0.2 (SI Fig. S5 E-F). In particular, the model predicts the slope of *Tn* vs *Tri* to be exactly one for all values of *x*, which is indeed observed experimentally (SI Fig. S5C). Altogether, the data and the model can be reconciled for the whole dataset if one assumes that the checkpoint is no more than 0.2*C* from the termination but most likely at the termination. It is important to emphasize that the agreement between the data and model can only be achieved for slow growth conditions (*Td* > 130 mins). For faster growth rates the model and the data do not agree for any values of *x*. This disagreement can be expected because the model assumes some replication-related process leading to onset of constriction. As argued above, this assumption is not likely valid at faster growth rates.

### Constriction can start before replication termination if the divisome is misplaced

Our goal was then to elucidate the molecular mechanism(s) that could be responsible for triggering constriction formation in a replication-dependent manner. Several molecular systems have been identified in the past that couple division and replication cycles in *E. coli* [32]. These include the nucleoid occlusion factor SlmA [33], the Ter linkage proteins ZapA, ZapB, and MatP [34, 35], and the DNA translocase FtsK [36]. The first two of these systems have been implicated in the positioning of the Z-ring relative to the replication terminus region of the chromosome while FtsK can reposition misplaced chromosomes relative to the division plane at the end of constriction [37]. We next asked if any of these systems are responsible for the correlated timing between the termination of replication and initiation of the constriction. We first considered the effects of SlmA, which is proposed to inhibit the formation of the Z-ring before the Ter region of the chromosome moves to the center of the cell in a replication-dependent manner [32, 38]. Its inhibitory effect is believed to be relieved from the mid-cell in the 2^nd^ half of the replication period because SlmA lacks binding sites at the Ter region. By removing SlmA from the cell one would expect the formation of the Z-ring and the constriction to start earlier. In contrast to this prediction, our data show that the *Tn* – *Trt* period increased compared to the WT strain in slow growth conditions (*p* = 4. 10^−4^; in single tailed Mann-Whitney test, Fig. 4A & Table S5). At the same time *R*(*Tn, Trt*) decreased compared to WT but still remained present at a significant level (*R* = 0.7; Fig. 4B). The observations clearly rule out the idea that SlmA is the main factor responsible for the timing of constriction formation. Rather, the above findings indicate that SlmA affects the timing of constriction indirectly by modulating the activity of some other factor.

**Figure 4.**
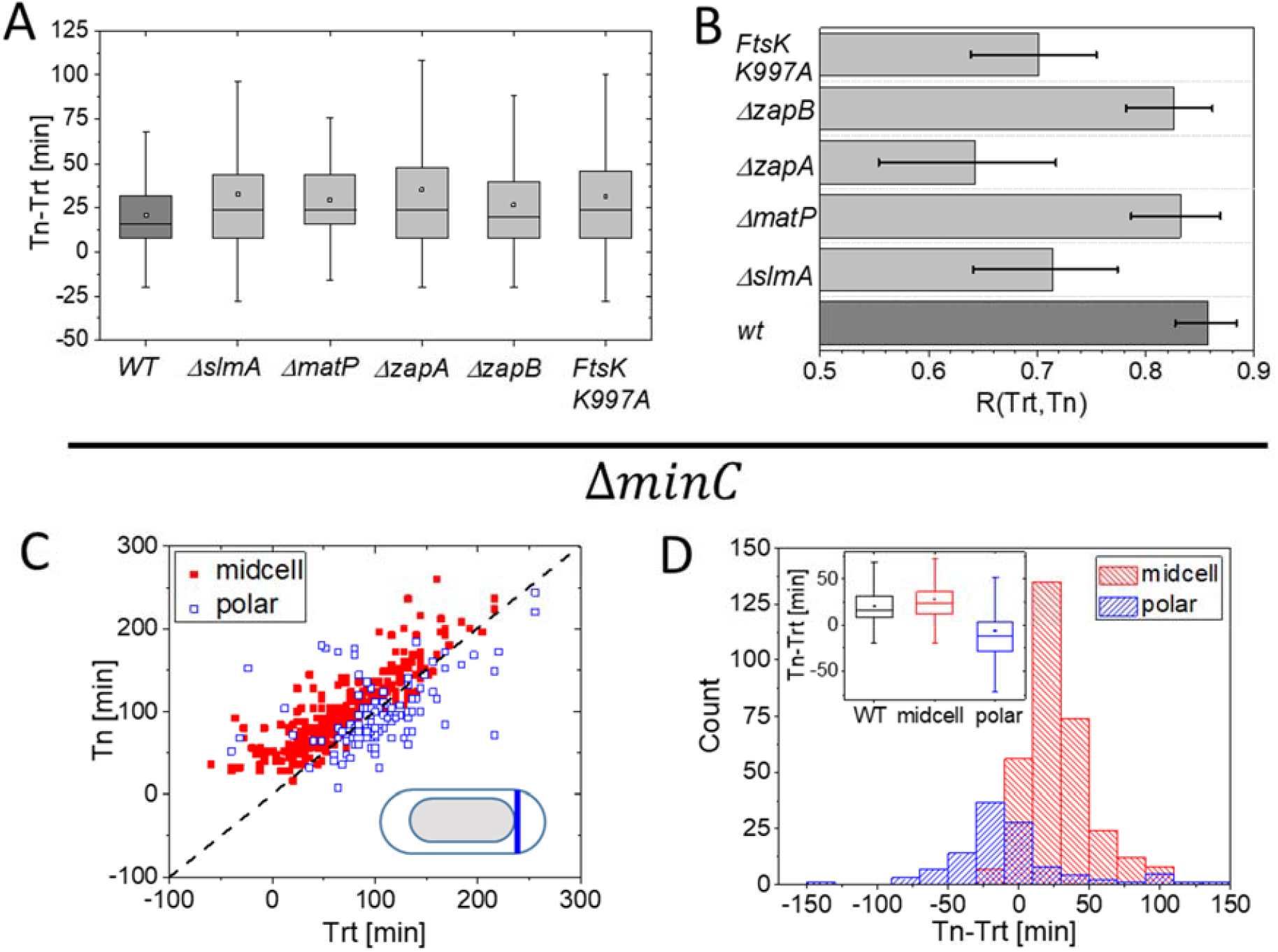
Timings for termination of replication and constriction initiation for different deletion mutants. The deleted gene products have been implicated in coordinating division and replication processes. (A) Delay times between the termination and initiation of constriction for wildtype (WT), Δ*slmA, ΔmatP, ΔzapA, ΔzapB*, and *ftsK K*997*A* strains. All mutant strains show longer delay times compared to WT strain at *p* = 0.05 level except Δ*zapA* (Mann-Whitney test; Table S5). (B) Pearson correlation coefficients between these times for the same strains. Error bars reflect 95% confidence intervals. (C) Termination of replication (*Trt*) vs initiation of constriction (*Tn*) for Δ*minC* cells. Polar divisions and mid-cell divisions are separately labeled. (D) Distribution of corresponding delay times in this strain. Inset compares the distributions to the corresponding one in WT cells. All measurements were performed in M9 Gly+TrEl medium.

Next, we investigated the role of the Ter linkage proteins ZapA, ZapB, and MatP. Unlike SlmA, which acts as an inhibitor, these proteins have been implicated in promoting the formation of the Z-ring [35]. Together ZapA, ZapB, and MatP form a proteinaceous chain that connects the replication terminus region of the chromosome to the Z-ring. For this connection, all three proteins are needed [34]. Since these proteins promote Z-ring formation, removal of either ZapA, ZapB, or MatP from cells should delay Z-ring formation and possibly also the formation of the constriction. Indeed, removal of either of these three proteins increased ⟨*Tn* – *Trt*⟩ in a statistically significant manner (SI Table S5) although the magnitude of the effect was small (less than 10% of cell cycle time; < 15 mins). The observed small increase in delay in constriction formation indicates that the Ter linkage proteins, similarly to SlmA, are unlikely to be directly involved in timing the constriction formation.

We also investigated the role of FtsK. FtsK has been implicated in segregating the replication terminus region at the onset of constriction [39, 40]. One could expect the unsegregated terminus region to delay constriction closure. Using a translocation defective mutant FtsK K997A [41] we indeed found ⟨*Tn* – *Trt*⟩ time to increase but the observed effect was again rather small (11 min; 7% of cell cycle time). Thus, all these mutants showed increased ⟨*Tn* – *Trt*⟩ periods compared to WT cells and slightly lower correlations in *R*(*Tn, Trt*) (Fig. 4B) but these effects were small enough to rule out their direct involvement in triggering the constriction formation. The observed small effects of these deletions arise likely via small changes these proteins have on the structure and composition of the divisome, and on the organization of the chromosome.

Although the Min system is not known to directly couple the replication and division processes, it is one of the main determinants for positioning the Z-ring in *E. coli* [32]. In cells with a defective Min system, a fraction of divisions occurs close to cell poles while the remaining ones still occur in the vicinity of cell middle. Distinguishing polar divisions from mid-cell ones shows that polar divisions can start significantly earlier than the mid-cell ones (Fig. 4C, D). About half of the polar divisions started before replication had terminated. At the same time, the timing of mid-cell divisions was not affected compared to WT cells (inset of Fig. 4D). The findings related to polar divisions rule out the possibility that termination *triggers* constriction formation as such a trigger would violate causality. On the other hand, these data raise the possibility that replicating nucleoids in mid-cell can block constriction formation. According to the previous discussion, this blockage is not dependent on the nucleoid occlusion factor SlmA.

## Discussion

We found that the initiation of the constriction and the termination of the replication in *E. coli* were poorly correlated at fast growth rates but the correlations increased as the growth rate slowed reaching *R* =0.94 for the slowest growth condition. The cross-over from a correlated to uncorrelated regime occurred approximately at *Td* ≈130 mins, which corresponds to *Td* ≈65 mins at 37 °C. A similar cross-over also appeared in *CV*(*Tn* – *Trt*) and in the slope of *Trt* vs *Tn* when plotted against *Td*. Furthermore, the distributions of delay times *Tn* – *Trt* become approximately exponential for *Td* > 130 mins suggesting that the replication is rate-limiting for the initiation of constriction.

One of our aims was to further elaborate on the coupling mechanism between replication and division cycles. The measurements with the *minC* deletion strain in slow growth conditions showed that in polar divisions initiation of constriction can precede termination of replication while the mid-cell divisions followed the same timings as in WT cells. This finding ruled out the possibility that termination acts as a trigger for constriction formation. Also, the finding ruled out that there can be a diffusible signal that triggers constriction formation which is released at [42] or before termination. The diffusible signal should reach within seconds all cellular locations including mid cell and pole, and it will not lead to observable differences in the timing of constriction formation. Note that a protein synthesized in response to transcriptional activation is a diffusible signal. The data thus rule out any possible mechanism where initiation of constriction is triggered in response to transcriptional activation of some gene. Instead of being triggered, the data from polar divisions suggested that a replicating and not fully segregated chromosome in the mid-cell blocks constriction formation; that is, replication related processes license the onset of constriction.

Although the coupling between the replication and division cycles appears to involve some form of nucleoid occlusion, it appears not directly related to the nucleoid occlusion factor SlmA. Clearly, there could be some unknown nucleoid occlusion factor that is not identified yet as argued before [35, 43, 44]. It is also possible that there are no additional proteins involved but the nucleoid occlusion arises directly from chromosome coils or transertion linkages [32], which present a steric hindrance for the formation and maturation of the Z-ring. These possibilities should be examined in further studies.

Altogether, our data indicate that there is a mechanism to license division in a replication-dependent manner (Fig. 5). In slow growth conditions, this licensing is rate-limiting for constriction formation. Our modeling studies suggest that the replication-dependent checkpoint occurs most likely at the termination but not earlier than 0.2*C* from the termination. Once the division is licensed, the constriction formation ensues via reaction kinetics which is suggestive of first-order reaction. At faster growth rates some other competing process appears to become rate-limiting. The origin of the “other” process remains to be also determined. Some authors have proposed the rate-limiting factors for the onset of constriction are precursor molecules for peptidoglycan synthesis [10] such as lipid II while others have concluded that it is the protein FtsZ [8]. Further work is thus needed to clarify the origin of this process.

**Figure 5.**
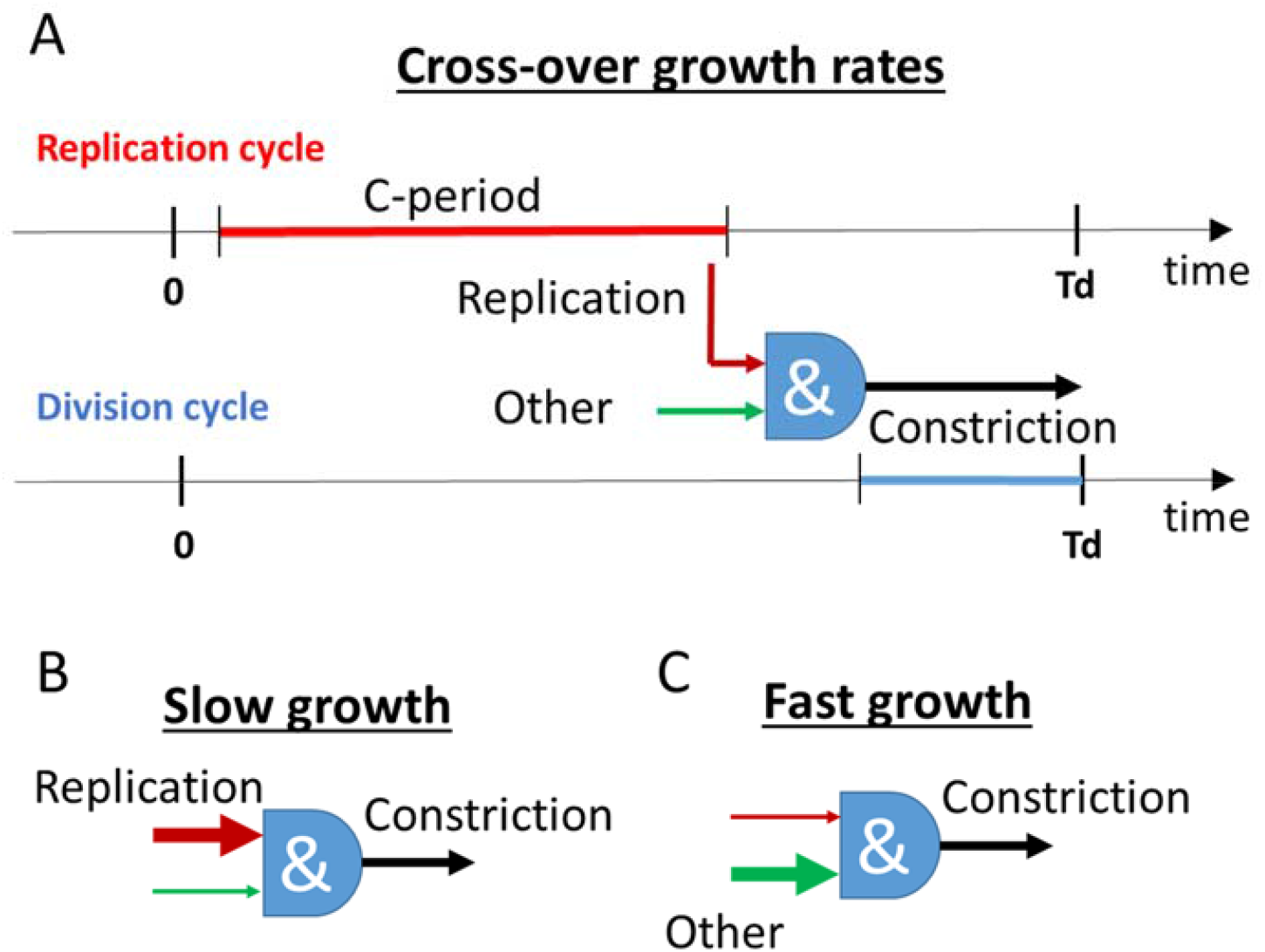
Regulation of constriction formation in different growth rates. (A) Regulation at the cross-over regime where *Td* ≈ 130 *min*. The corresponding doubling times at 37 °C are expected to be about twice shorter. The replication period is shown by red and the constriction period by blue lines. & sign indicates an integration of different signals. Constriction starts when conditions imposed by the replication and by some “other” yet to be identified processes have been both met. Replication related processes relieve inhibition for initiation of constriction at or shortly before the termination. (B) In slow growth conditions, the onset of constriction is rate-limited by replication-related processes. (C) In fast growth conditions, some unknown “other” process(es) become rate-limiting.

The regulation proposed in Fig. 5 is similar to the concurrent processes model [11, 12] with some differences. First, the concurrent processes model does not consider the initiation of the constriction as a cell cycle checkpoint. Instead, it predicts the timing and cell size at the division; that is at the end of all division-related processes. Note that all other current cell cycle models in *E. coli* also predict only the end of the division. Second, in the concurrent processes model replication and growth-related processes are both rate-limiting for division in all growth conditions. Our data suggest that replication is rate-limiting for constriction only in slow growth conditions and the concurrency of the processes is only significant at the vicinity of the cross-over region.

In conclusion, our work has shown that cell division is limited by replication-related processes in slow growth conditions but appears to be almost independent of these processes in fast growth. This behavior might explain why some earlier works have inferred that replication and division cycles are completely uncoupled from each other [8–10] while other authors have come to exactly opposite conclusions [1, 2, 6, 7]. Our data furthermore implies that the limitation related to replication processes stems from some yet to be identified form of nucleoid occlusion. This nucleoid occlusion is lifted in a replication-dependent manner. In parallel to this limitation, cells experience other types of licensing conditions that need to be met. Whether all these limiting processes couple to the divisome via the central hub of FtsZ protofilaments or also via other divisome components remains to be elucidated.

## Acknowledgments

The authors thank Ethan Garner, Conrad Woldringh and Arieh Zaritsky for useful discussions, Da Yang and Scott Retterer for help in microfluidic chip making. Authors acknowledge technical assistance and material support from the Center for Environmental Biotechnology at the University of Tennessee. A part of this research was conducted at the Center for Nanophase Materials Sciences, which is sponsored at Oak Ridge National Laboratory by the Scientific User Facilities Division, Office of Basic Energy Sciences, U.S. Department of Energy. This work has been supported by the US-Israel BSF research grant 2017004 (JM), the National Institutes of Health award under R01GM127413 (JM), NSF CAREER 1752024 (AA) and NSF award 1806818 (PK).

## STAR METHODS

### KEY RESOURCES TABLE

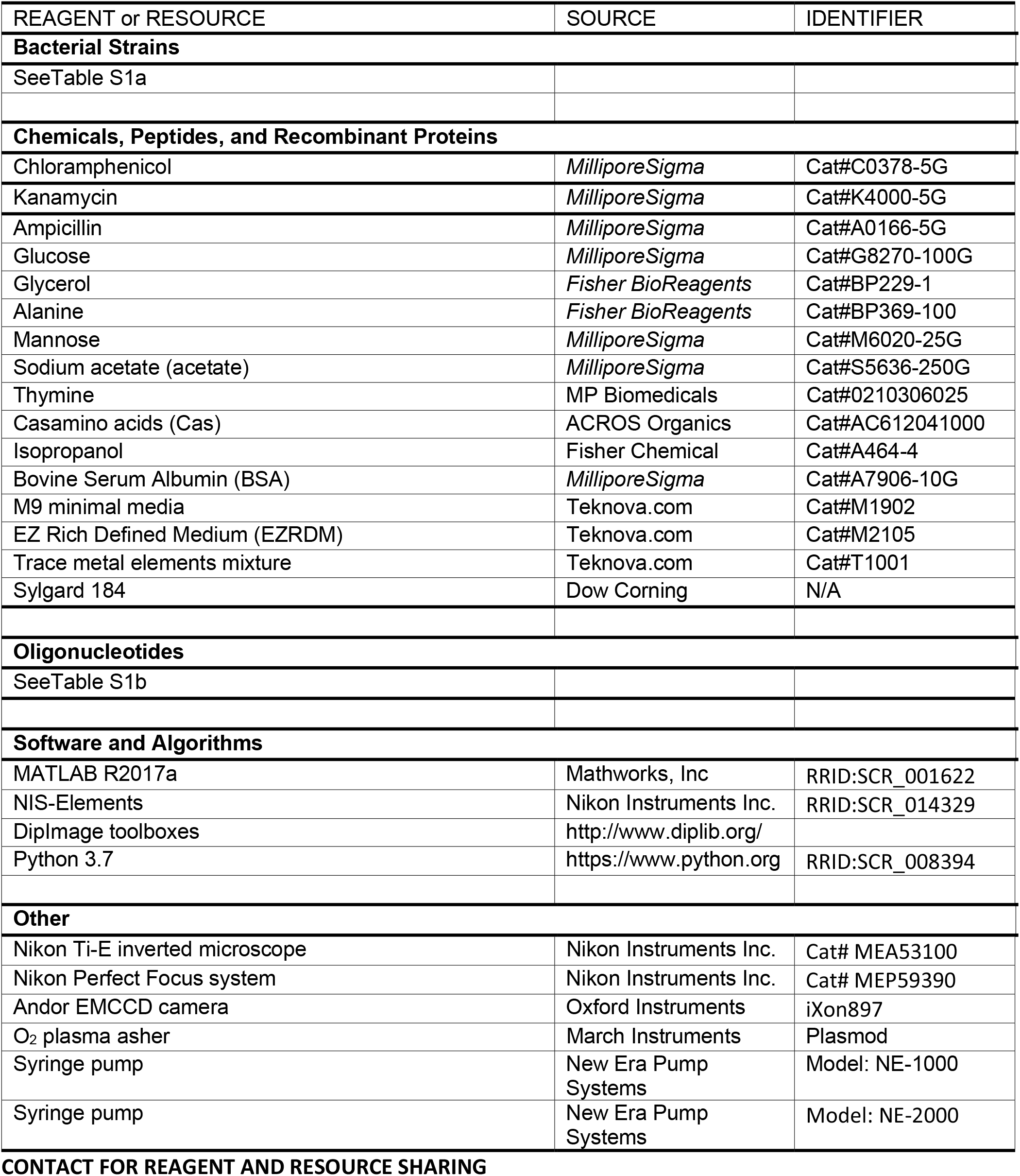

### CONTACT FOR REAGENT AND RESOURCE SHARING

Further information and requests for resources and reagents should be directed to and will be fulfilled by the Lead Contact, Jaan Männik.

## METHODS

### Construction of *E. coli* strains

All *E. coli* strains used in the reported experiments are derivatives of K12 BW27783 obtained from the Yale Coli Genetic Stock Center (CGSC#: 12119). Strains were constructed either by λ-Red engineering [45] and/or by P1 transduction. Where necessary kanamycin resistance gene was removed by expressing the Flp recombinase from plasmid pCP20 [46]. Detailed information of strain genotypes and construction information is listed in Table S1a. Oligonucleotide information is given in Table S1b. For *E. coli* strain engineering, cells were grown in lysogeny broth (LB) and appropriate selective antibiotics.

### Growth media and growth conditions

For time-lapse imaging in microfluidic devices, cells were cultured in 9 different growth conditions at 28°C. Detailed information on the media used can be found in SI Table S3.

### Cell preparation and culture in microfluidic devices

All bacterial strains were streaked on agar plates containing M9 minimal salts supplemented with 2 mM magnesium sulfate, corresponding carbon sources, and appropriate selective antibiotics. A day before an experiment a less than 10 days old colony was inoculated into 3 ml of EZ Rich Defined Medium (EZRDM, Teknova Inc., CA) or M9 minimal salts media (Teknova Inc., CA) supplemented with corresponding carbon sources, trace metals mixture (Teknova Inc., CA, #T1001), casamino acids (ACROS Organics) and appropriate antibiotics when needed. Unless otherwise indicated, antibiotics were used at 25 µg/ml of kanamycin (Kan) and 25 µg/ml chloramphenicol (CM). For microscopy experiments, cells were grown to an OD_600_ of ∼0.1 in a liquid medium and then concentrated ∼100x by centrifugation in presence of 0.75 µg/ml of BSA (Bovine Serum Albumin; Millipore Sigma, MO) to minimize clumping of the cells. The resulting solution was used to inoculate microfluidic mother machine devices. The latter were made of PDMS (polydimethylsiloxane) following a previously described procedure [30]. For inoculation 2-3 µl of resuspended concentrated culture was pipetted into the main flow channel of the device. The cells were then let to populate the dead-end channels. Once these channels were sufficiently populated (about 1 hr), tubing was connected to the device, and the flow of fresh M9 medium with corresponding carbon sources and supplements, and BSA (0.75 µg/ml) was started. The flow was maintained by a NE-1000 Syringe Pump (New Era Pump Systems, NY) at 5 µl/min during the entire experiment. To ensure steady-state growth, the cells were left to grow in channels at least 14 hr (24 hrs for acetate) before imaging started.

### Fluorescence microscopy

A Nikon Ti-E inverted fluorescence microscope (Nikon Instruments, Japan) with a 100X NA 1.40 oil immersion phase contrast objective (Nikon Instruments, Japan), was used for imaging the bacteria. Images were captured on an iXon DU897 EMCCD camera (Andor Technology, Ireland) and recorded using NIS-Elements software (Nikon Instruments, Japan). Fluorophores were excited by a 200W Hg lamp through an ND4 and ND8 neutral density filter. Chroma 41004 and 41001 filter cubes (Chroma Technology Corp., VT) were used to record mCherry and Ypet images, respectively. A motorized stage (Prior Scientific Inc., MA) and a Nikon Perfect Focus ® system were utilized throughout time-lapse imaging.

### Image analysis

MATLAB, along with the Image Analysis Toolbox and DipImage Toolbox, (http://www.diplib.org/) were used for image analysis. In all analyses of time-lapse recordings, corrections to subpixel shifts between different frames were applied first. These shifts were determined by correlating phase-contrast images in adjacent frames. The cells were then segmented based on phase-contrast images using a custom MATLAB script. The segmented images were used to compile heatmaps of phase and fluorescent images as shown in Fig. 1B. Timings of cell divisions were corrected based on the dissociation of the Ypet-FtsN label from the septum in strains where this label was present. Timings of replication initiation and termination, and initiation of constriction were determined from heatmaps. For the Figures in the main text the replication initiation, *Tri*, and termination, *Trt*, timings were determined from the heatmaps of the mCherry-DnaN label (strain STK13). For a detailed procedure see below. In glycerol and glucose+Cas growth media, these timings were also determined using a different strain (JM85), which expressed the ssb-Ypet label. The data from the latter strain are shown in SI Figures. Timings for the onset of FtsN recruitment to the Z-ring, *Tn*, were determined from Ypet-FtsN heatmaps. The onset of constriction, *Tc*, was determined from phase hetamaps for both strains.

### Determination of *Trt* timing

In slow growth conditions, there is typically a single termination that increases the number of chromosomes from one to two while at faster growth rates a fraction of cells is born with two chromosomes. To analyze cells born with one and two chromosomes on the same footing we considered the relevant termination for the cells born with two chromosomes to occur in the mother cell. Using this analysis, the termination times of the cell population have a continuous unimodal rather than bimodal distribution. In this distribution, the times of replication termination are negative if these times occur in the mother cell. The two terminations in the mother cell were not exactly synchronous. For a given cell of interest, we determined the timing of the termination for the chromosome that was inherited by this cell.

### Determination of *Tri* timing

Similar to the termination, the relevant initiations of replication could occur in the mother cell. In the two fastest growth rates, in EZ-Rich and M9 glucose+CAS the relevant initiations could also occur in the grandmother cells. Our analysis routine did not allow us to determine these events. Also, in these two growth conditions even when the initiations occurred in the mother cell it was rather ambiguous to determine their timing. We, therefore, left out these two growth conditions from the analysis that involved replication initiation (in SI Fig. S5 and Table S4). The timing of replication termination (*Trt*) could still be reliable determined in these conditions. In other growth media, the initiation occurred either in the mother cell or in the cell of interest. If the initiation occurred in the mother cell then the timing of the initiation of the chromosome that was inherited by the cell of interest was used. The time difference between the two initiations in the mother cell was typically within 8 min interval.

### Statistical Analysis/Error Analysis

Different distributions of *Tn* – *Trt* times were compared using t-test and Mann-Whitney test. For testing Matlab functions *ttest2* and *ranksum* were used, respectively.

Error bars for the Pearson R-values represent 95% confidence intervals. These intervals were calculated using Fisher’s z-transformation [47]. Briefly, based on the measured R-value corresponding z-value was calculated as *z* = 1/2 *ln*(1 + *R*/1 – *R*). The 95% confidence intervals for z were calculated as 3, *z* + 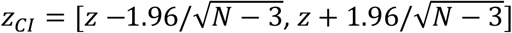 where *N* is number of measurements for a given R-value. The intervals were then backtransformed for R confidence intervals using *R*_CI_ = (*exp*(2*z_CI_*) – 1)/(*exp*(2*z_CI_*) + 1).

Error bars for the coefficient of variation (CV) also represent 95% confidence intervals. These intervals were determined by bootstrapping. Bootstrapping was carried out in Phyton 3.7. by sampling the distributions 10^4^ times and verifying that the resulting CV distributions did not change upon further doubling the samples. Percentile intervals were found using *numpy percentile* method.

### Model coupling replication and constriction

We assume that cells initiate DNA replication at time *Tri* and that nucleotides are added to the growing strand of DNA at a constant rate *γ*. We denote the length of the *E. coli* genome (measured in nucleotide number) as *N*. Using the central limit theorem, the time taken to reach termination (the C period) is normally distributed with a mean 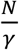 and variance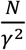. The CV of the *C* period thus scales as 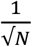 with *N* ≈ 4. 10^6^. Thus, the predicted CV is two orders of magnitude smaller than that experimentally observed (SI Table S4), and we conclude that the variability in the *C* period resulting from stochastic nucleotide addition is negligible. Hence, we can consider the replication process to be happening at a constant velocity *v*. The time taken for replication to complete is then given by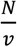, and variability in the *C* period results from the cell-to-cell variability in *v*.

Experimentally, the *C* period and *Td* are strongly and positively correlated (SI Table S4). This would suggest that the progress of biochemical processes like DNA replication scales with the individual growth rate of the cells. In this case, a slower-growing cell will also replicate at a slower velocity and subsequently have a longer *C* period. The scaling with growth rate points to a small but non-negligible variability in *v* within the population of growing cells in a particular media.

Since *C* is assumed uncorrelated with the initiation time, the time at termination *Trt*′ is thus:

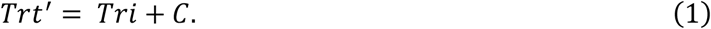

The evidence for such a “timer” can be found in SI Table S4, which lists the slope of linear regression for *Trt* vs *Tri* plots in different growth conditions. As can be seen from SI Table S4, this slope is close to one in slow growth conditions.

In the experiments, the replisome is imaged using a DnaN marker. The DnaN marker is expected to remain attached to the replication terminus region after completion of replication [25]. Thus, in experiments the measured time of termination *Trt* is,

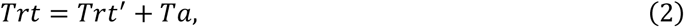

where *Ta* is the time for which DnaN stays attached. *Ta* is assumed to be exponentially distributed with a mean time ⟨*Ta*⟩ = 3-6 mins expected in our growth conditions [25].

We assume *Tri* to be normally distributed with mean ⟨*Tri*⟩ and standard deviation σ_ri_, the values for which are determined from experiments. Assuming *v* to be normally distributed with a mean *v*_O_ and standard deviation σ_v_, the *C* period has an approximately normal distribution with mean 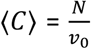 and variance 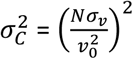 when σ ≪ *v* _0_. Using Equations 1 and 2, we can determine ⟨*C*⟩ and 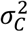 to be,

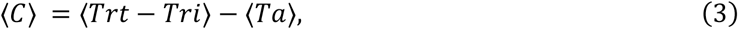

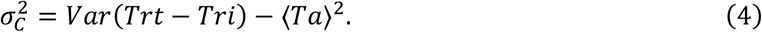

⟨*Trt* – *Tri*⟩ and *Var*(*Trt* – *Tri*) are the mean and variance of *Trt* – *Tri* and are determined directly from experiments (SI Table S4).

In our model, constriction is said to be controlled by an event placed at a locus that is a relative distance of *x* from the replication terminus. *x* = 0 denotes the locus is at the terminus while *x* = 1 denotes a checkpoint at the initiation. Under the assumption that *v* has a Gaussian distribution with CV≪ 1, we obtain that the checkpoint is triggered after a time *ξ* from initiation which is normally distributed with mean 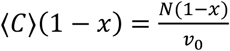 and variance 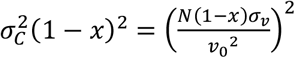. Thus, the checkpoint is said to be reached at time *Tx* given by,

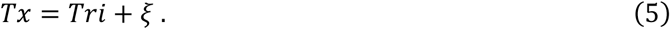

Since termination happens at a fraction *x* along the genome from *Tx*,

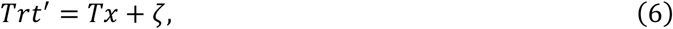

where *ζ* is normally distributed with mean ⟨*C*⟩*x* and variance 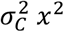. Note that *ξ* and *ζ* are correlated with each other with covariance Cov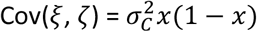. Both *ξ* and *ζ* are also correlated with the *C* period.

Constriction is assumed to be triggered by the checkpoint at time *Tx* at a constant rate *r*. This is based on the fact that the positive values of *Tn* – *Trt* are exponentially distributed (Fig. 1D) and the CV of *Tn* – *Trt* is close to one (Fig. 2D). Hence, the time at constriction *Tn* is,

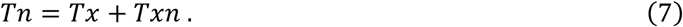

*Txn* is exponentially distributed with a mean time = 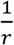. Using eqs. 2 and 6, we get

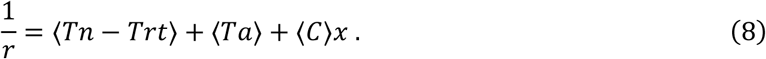

⟨*Tn* – *Trt*⟩ can be determined from experiments thus fixing the rate *r* for different *x. x* is a free parameter whose value is yet to be determined. Using the experimental results plotted in Fig. 2 and comparing them against the analytical results for varying *x*, we obtain constraints on the value of *x*.

We shall first calculate analytically CV(*Tn* – *Trt*) for a given value of *x*, which can be compared to experimentally determined values shown in Fig. 2D. Using eqs. 2, 6, and 7, we get,

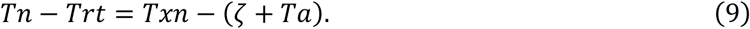

*Txn, ζ, Ta* are independent of each other. Hence the variance of *Tn* – *Trt* is

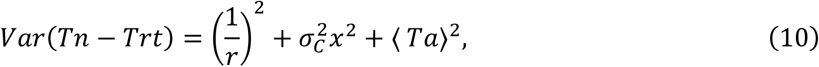

while the mean is 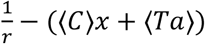. Combining the two, we find:

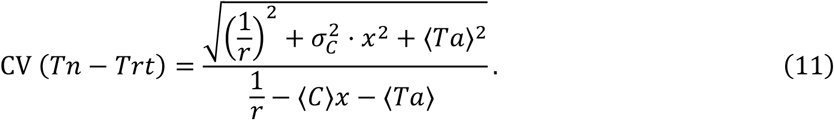

Other statistical constructs presented in Fig. 2 include the Pearson correlation coefficient for *Trt, Tn, R*(*Tn, Trt*) and the slope of linear regression for *Tn* vs *Trt* plot.

*R*(*Tn, Trt*) is defined as,

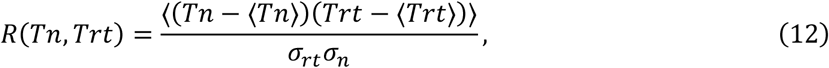

where σ_rt_ and σ_n_ are the standard deviations of *Trt* and *Tn*, respectively.

From eqs. 5 and 7, we obtain *Tn* = *Tri* + *Txn* + *ξ*. Similarly, from eqs. 1 and 2, *Trt* = *Tri* + *C* + *Ta*. All pairs of variables from *Tri, C, Ta, Txn, ξ* are uncorrelated with each other except *C* and *ξ* which are correlated as remarked earlier. Substituting this into eq. 12, we find the covariance (numerator) to be,

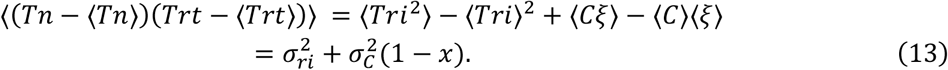

σ_n_ and σ_rt_ are found to be:

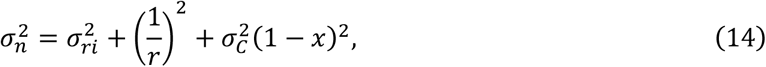

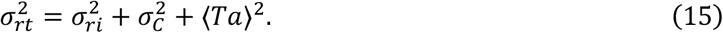

Substituting eqs. 13, 14, and 15 into eq. 12, we can obtain *R*(*Tn, Trt*). All the parameters in the formula for *R*(*Tn, Trt*) can be extracted from experiments while *x* is a variable.

The slope of the linear regression line for *Tn* vs *Trt* is related to *R*(*Tn, Trt*) as

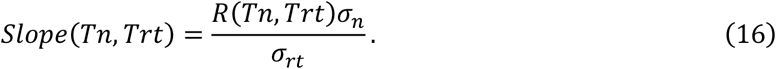

This can also be calculated by substituting the values in eq. 13, 14, and 15. This theoretical prediction is compared to the experimental data in Fig. 2C.

Assuming the trigger for the constriction event to be at termination (i.e., *x* = 0), we can obtain the distribution of *Tn* – *Trt* times analytically and compare it to experimental distributions. For *x* = 0, we obtain *Tn* – *Trt* = *Txn* – *Ta*. Let us define the random variable *Z* = *Tn* – *Trt*. We aim to find its distribution. Using our assumptions that *Txn* and *Ta* are independent and exponentially distributed, we obtain the joint probability distribution of *Txn* and *Ta* to be,

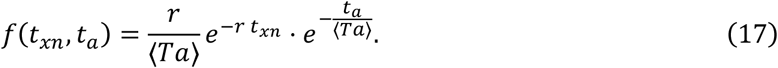

For *z* ≥ 0,

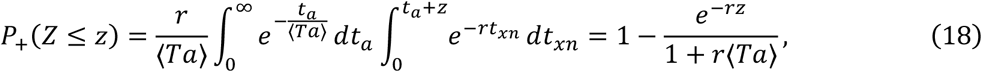

with *P*_+_(*Z* ≤ *z*) the cumulative distribution function (CDF) of *Z* for *z* ≥ 0. Similarly, for *z* ≤ 0, we obtain,

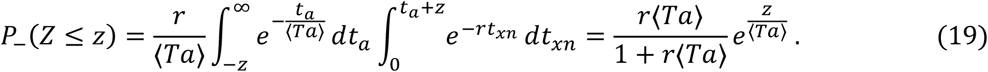

Therefore, we find that the probability distribution of 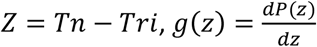 is

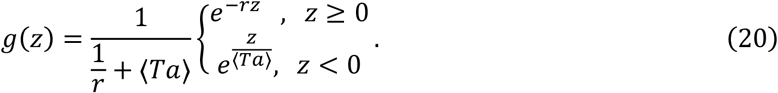

The parameters can be determined using the experimental data as discussed before.

Finally, we also investigate the relationship between replication initiation timing *Tri* and timing for initiation of constriction *Tn*. As before we will calculate the relevant statistics as a function of *x*. We will rely on the fact that *Tn* = *Tri* + *Txn* + *ξ*, and that *Tri, Txn* and *ξ* are uncorrelated. The variance of *Tn* – *Tri* is found to be,

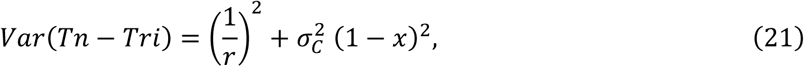

while the mean of 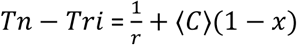. Thus, CV(*Tn* – *Tri*) is found to be,

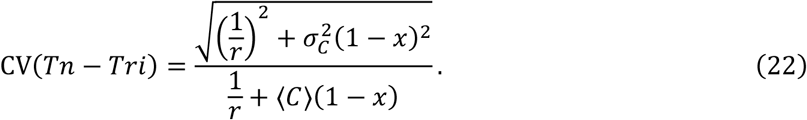

Other statistical constructs which we calculate include the Pearson correlation coefficient for *Tri, Tn, R*(*Tn, Tri*) and the slope of linear regression for *Tn* vs *Tri* plot.

*R*(*Tn, Tri*) is defined as,

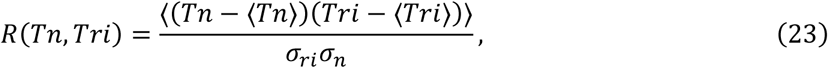

Using *Tn* = *Tri* + *Txn* + *ξ*, we find the numerator to be:

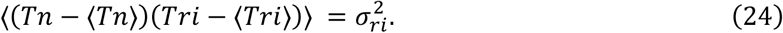

The quantity σ_ri_ is directly inferred from experiments while σ_n_ is calculated using eq. 14. Substituting the values into eq. 23, we can obtain *R*(*Tn, Tri*). The slope of the linear regression line between *Tn* and *Tri* is related to *R*(*Tn, Tri*) as

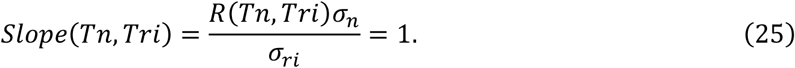

Hence, the slope of the linear regression line for *Tn* vs *Tri* is always one independent of growth conditions. In other words, within the model *Tn* is related to *Tri* via a timer. In the four slowest growing conditions, the slope between *Tn* vs *Tri* is indeed close to 1 as shown in SI Fig. S5C.

