## Supplemental Figures and Tables for "Replication-related control over cell division in *Escherichia coli* is growth-rate dependent"

Supporting Information contains:

1) Figures S1-S5

2) Tables S1-S5

3) References

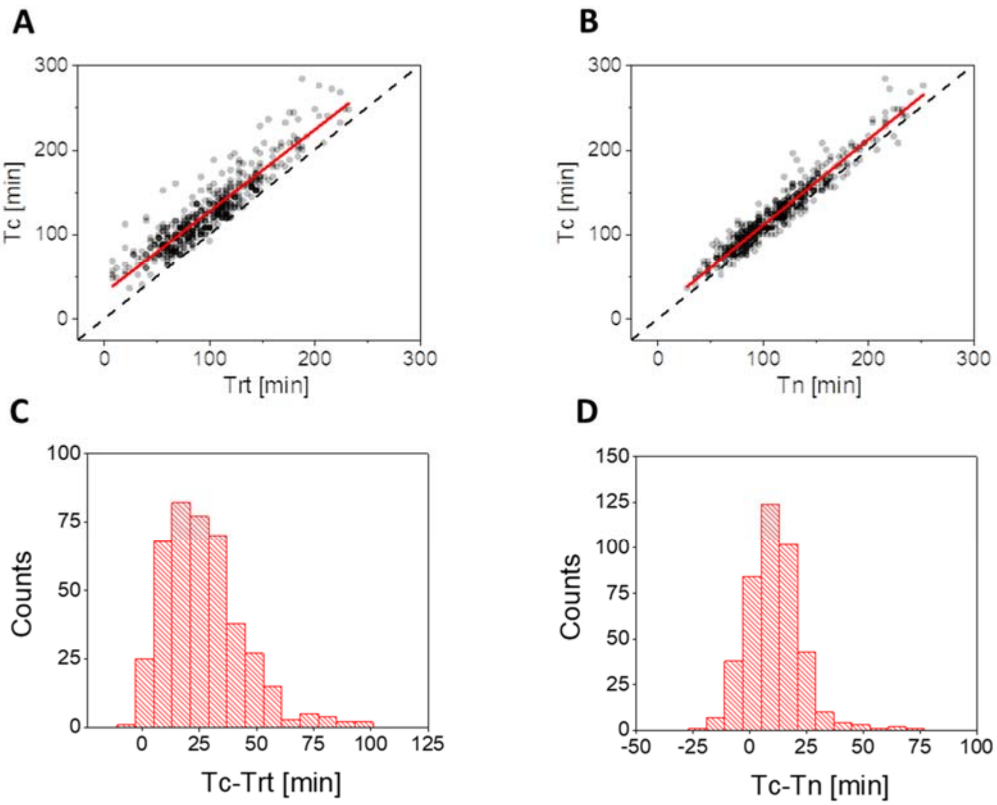


**Figure S1.** (A) Correlations between the termination of replication ($Trt$) and initiation of constriction ($Tc$). Timing for $Tc$ is determined from the phase images. The solid red line is linear fit ($Tc = 0.97Trt + 31$; $R = 0.92$) and the dashed black line corresponds to $Trt = Tc$. The dashed black line corresponds to $Tc = Trt$. (B) Correlations between recruitment time of FtsN to mid-cell ($Tn$) and $Tc$. The solid red line is a linear fit ($Tc = 1.01Tn - 10$; $R = 0.96$) and the dashed line $Tc = Trt$. (C) Distribution of delay times between $Trt$ and $Tc$ ($28 \pm 18$ mins; $mean \pm std$). (D) Distribution of delay times between $Tn$ and $Tc$ ($12 \pm 12$ mins; $mean \pm std$). $N = 420$. All measurements were taken in a glycerol medium. Strain STK13.

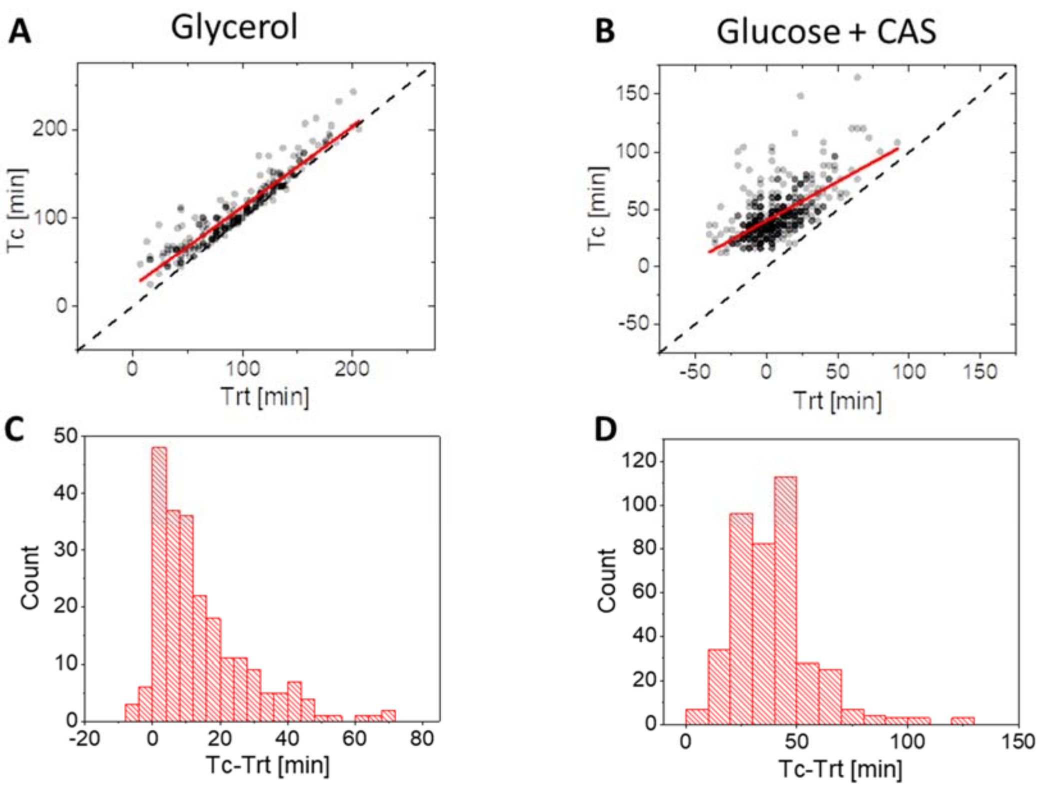


**Figure S2.** Measurements in strain JM85 carrying ssb-mYpet label for replisome and no divisome label. The timing for constriction initiation ($Tc$), is determined from the phase images. (A) Correlations between the termination of replication ($Trt$) and initiation of constriction ($Tc$) in slow growth conditions in glycerol medium (without TrEl). The solid red line is linear fit ($Tc = 0.91Trt + 23; R = 0.94$) and the dashed black line corresponds to $Trt = Tc$. (B) Correlations between $Trt$ and $Tc$ in the fast-growth condition in Glucose+Cas medium. The solid red line is linear fit ($Tc = 0.68Trt + 40; R = 0.62$) and the dashed line is the same as in (A). (C) Distribution of delay times between $Trt$ and $Tc$ for these cells in slow-growth conditions ($21 \pm 18$ mins; mean $\pm$ std). (D) Distribution of delay times between $Trt$ and $Tc$ for these cells in fast-growth conditions in glucose+Cas medium ($38 \pm 18$ mins; mean $\pm$ std).

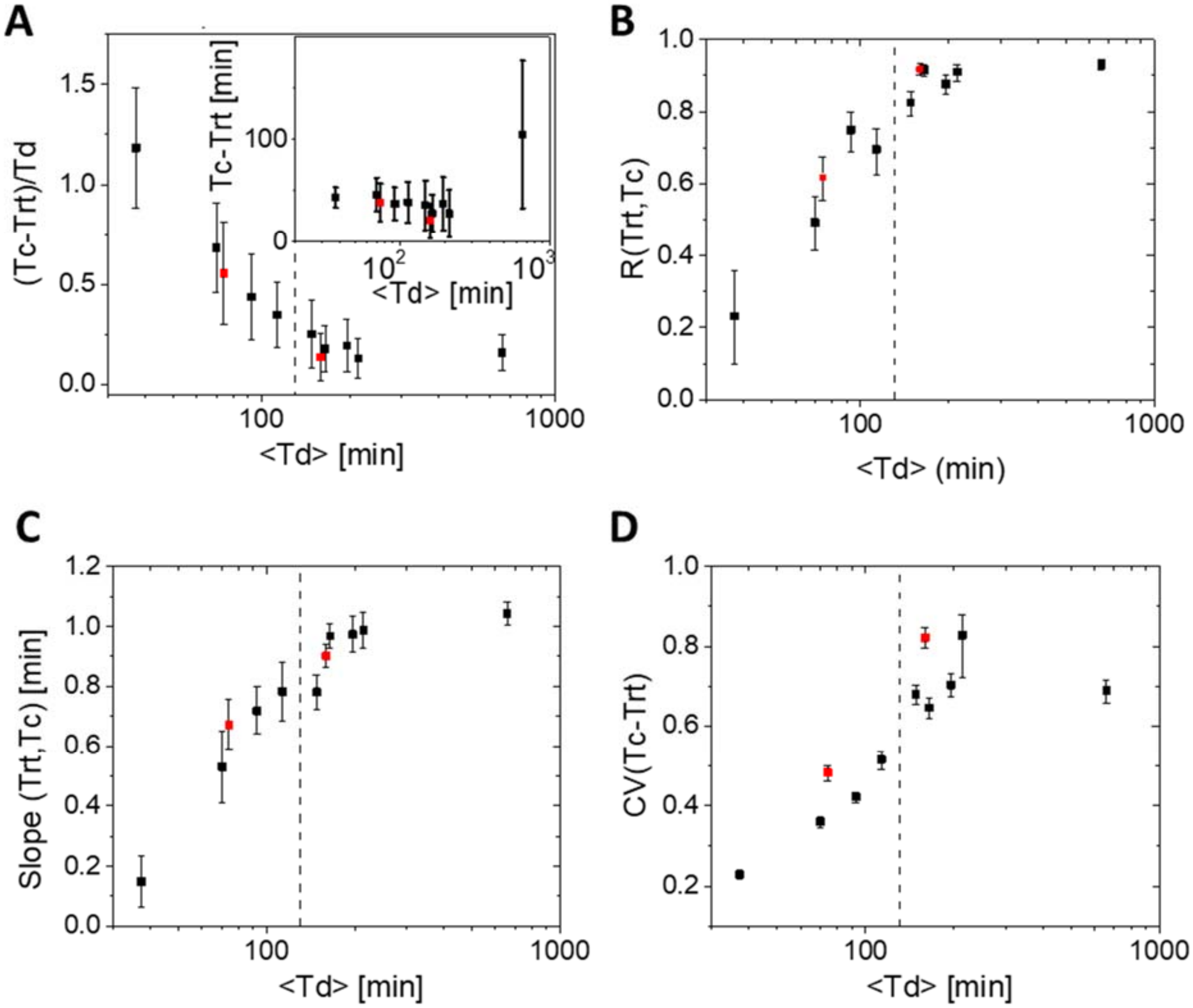


**Figure S3.** Timings of constriction initiation determined from the *phase images*, $Tc$, relative to replication termination, $Trt$, in 9 different growth conditions (listed in Table S3). Black points correspond to strain STK13, which carries mCherry-DnaN and Ypet-FtsN. Red points correspond to strain JM85, which carries an only ssb-mYpet label. The Figure here follows the same layout Fig. 2 in the main text but $Tc$ replacing $Tn$ here. (A) The normalized average delay time between initiation of constriction and termination of replication as a function of the average doubling time, $< Td >$. The inset shows the same plot for the actual delay times. Error bars in both plots are std of these quantities within the cell population. (C) Pearson correlation coefficient between $Trt$ and $Tc$. (D) The slope of $Trt$ vs $Tc$ plot. (E) Coefficient of variation for $Trt - Tn$ distribution. The dashed vertical lines in all plots correspond to $< Td > =$ $130$ min. Error bars in (C)-(E) correspond to 95% confidence intervals. For calculation of these intervals see SI Text.

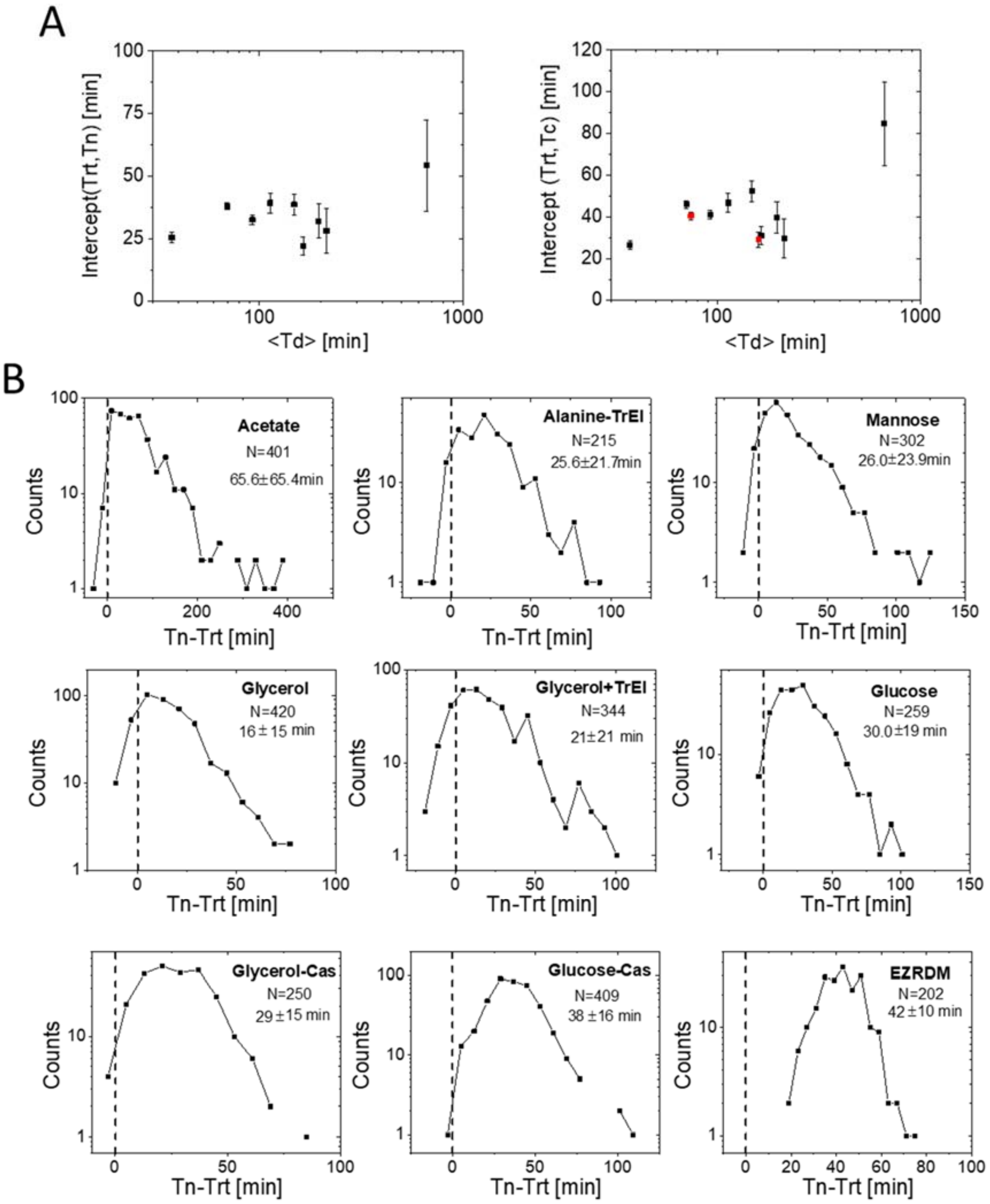


**Figure S4.** (A) Intercepts from linear fits to $Trt$ vs $Tn$ data (left panel) and to $Trt$ vs $Tc$ data (right panel) as a function of the average doubling time. Black points are from strain STK13 and red points from strain JM85. Error bars show 95% confidence intervals. (B) Distribution of $Tn - Trt$ times in the 9 different growth conditions arranged in the descending order of doubling times from the slowest growth condition at top-left to the fastest growth condition at bottom-right. $N$ is the number of analyzed cells and the numbers underneath it show the mean ± std for the distribution. Note that the y-axes are logarithmic.

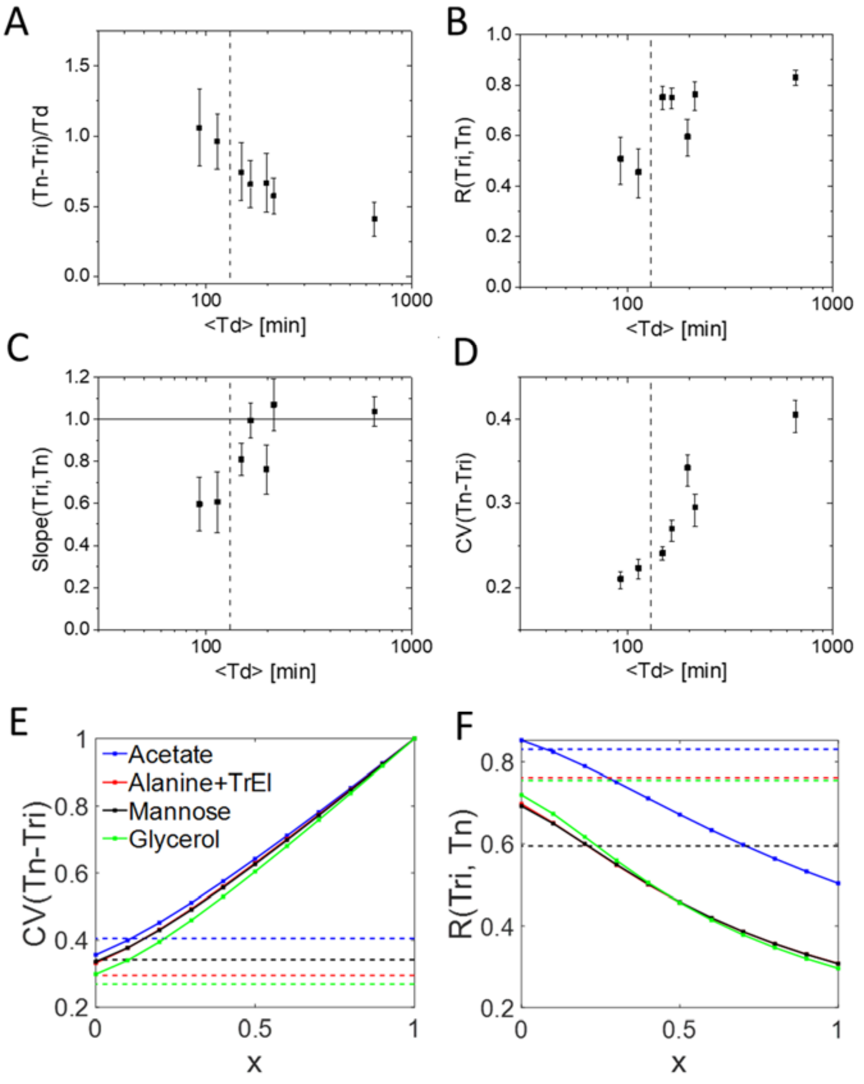


**Figure S5:** Statistics for the determined timings of constriction initiation, $Tn$, and initiation of replication, $Tri$, in 7 different growth conditions (panels A-D), and comparison of these statistics to the model (panels E-F). From the longest to shortest doubling times the carbon sources used in the media are acetate, alanine, mannose, glycerol, glycerol + trace elements (TrEl), glucose, glycerol+Cas (for details see Table S3). Note that the two fastest growth conditions reported in the main text are not shown here because the determination of the initiation times for DNA replication in these conditions becomes ambiguous. (A) The average normalized delay time between initiation of constriction and initiation of the replication as a function of the average doubling time, $< Td >$. Error bars in both plots show the std of these quantities within the cell population. (B) Pearson correlation coefficient between $Tri$ and $Tn$. (C) The slope of $Tri$ vs $Tn$ plot. A solid black line shows the prediction of the model. The same theoretical prediction

($slope(Tri, Tn) = 1$) applies for all growth conditions. (D) Coefficient of variation for $Tri - Tn$ distribution. The dashed vertical lines in all plots correspond to $< Td >= 130$ min. Error bars in (C)-(F) show 95% confidence intervals. (E) Analytical results for the coefficient of variation for the $Tn - Tri$ distribution as a function of $x$. (F) The same for the Pearson correlation coefficient between $Tri$ and $Tn$. In panels (E)-(F) the solid lines show predictions of the model. The dashed horizontal lines indicate the mean experimental values. Note that only the four slowest growth conditions are considered in these comparisons and $\langle Ta \rangle = 3\ min$. For details of calculations, see Methods, Model.

**Table S1a.** List of the strains constructed and used in the study.

| Strain | Genotype | Notes |
|---|---|---|
| BW27783 | Δ(araD-araB)567, ΔlacZ4787(::rrnB-3), λ-, Δ(araH-araF)570(::FRT), ΔaraEp-532::FRT, ϕ$P_{cp8}$araE535, rph-1, Δ(rhaD-rhaB)568, hsdR514 | CGSC Strain#: 12119 |
| STK13 | BW27783 ΔftsN::frt-Ypet-ftsN, ΔdnaN::frt-mCherry-dnaN | Ypet-ftsN: λ-Red, frt-*kan$^{R}$-frt-Ypet-linker*[a], pCP20, PCR; mCherry-dnaN: P1 transduction from BN1682[b], pCP20 |
| JM85 | BW27783 Δssb::ssb-Ypet-frt-kan-frt | P1 transduction from RRL32[c] |
| JM111 | BW27783 STK13 ΔftsK::ftsK$^{K997A}$-CM | P1 transduction from FC1[d], PCR, sequencing |
| JM116 | BW27783 STK13 ΔminC::frt-kan-frt | P1 from JW1165 (CGSC#: 9075), PCR |
| JM121 | BW27783 STK13 ΔslmA::frt-kan-frt | P1 from *JW5641-1 (CGSC#: 11495*), PCR |
| JM122 | BW27783 STK13 ΔzapA::frt-kan-fr | P1 from JW2878-1 (CGSC#: 10232), PCR |
| JM123 | BW27783 STK13 ΔmatP::frt-kan-frt | P1 from JW0939-1 (CGSC#: 12061), PCR |
| JM127 | BW27783 STK13 ΔzapB::frt-kan-frt | P1 from *JW3899-1* (CGSC#: 10814), PCR |

[a] a kind gift from R. Reyes-Lamothe, McGill University, Canada; [1]
[b] a kind gift from N. Dekker, TU Delft, The Netherlands; [2]
[c] a kind gift from R. Reyes-Lamothe, McGill University, Canada; [3]
[d] a kind gift from F. X. Barre, CNRS, France; [4]

**Table S1b**. List of oligos used in the study.

| Oligo | Sequence 5’>3’ | Notes |
|---|---|---|
| ftsN-F | CATGGCGGGCTGACGAACGAATAAATACAG CGAAACGATATGTAGGCTGGAGCTGCTTCG | λ-Red > Ypet-ftsN |
| ftsN-R | AAGGTGCCGGTTGGCTGCGGCGTACATAAT CTCGTTGTGCCGCGCTGCCAGAACCAGCGG | λ-Red > Ypet-ftsN |
| ftsN-test-F | TTACGGCGTACGCTTACTAT | verification of Ypet-ftsN |
| ftsN-test-R | CTTTAAGACTGCCGTGACTG | verification of Ypet-ftsN |
| minC-left | CAGCGCCATTTATCACAGAA | verification of ΔminC |
| minC-right | CCTTTGCCCGAAGTAACAAC | verification of ΔminC |
| matP-left | TTCACTGGCCTAAAAAGCTGA | verification of ΔmatP |
| matP-right | CGCAGGCTTAAGTTCTCGTC | verification of ΔmatP |
| slmA-left | TTGGTCACTCTGGTCGTCAG | verification of ΔslmA |
| slmA-right | AGGCCTATCGCGAAGAGTTT | verification of ΔslmA |
| zapA-left | GCAGTCAATCAGCAGGAAGG | verification of ΔzapA |
| zapA-right | CTTGCGAACATCTCAGAGAAA | verification of ΔzapA |
| zapB-F | GTAATCGGGACGAGGATTT | verification of ΔzapB |
| zapB-R | TTCTGCGTTACCTGTTGG | verification of ΔzapB |
| ftsK-F | GTGCGTGTCGTTGAAGTTATTC | verification of FtsK $^{K997A}$ |
| ftsK-R | GTTTATACCGACGCTCCATCTC | verification of FtsK $^{K997A}$ |

**Table S2:** Cell characteristics data of strains used in the study (mean ± SD). $L_b$ corresponds to cell length at birth.

| Strain | Media | No. of cells | Td [min] | $L_b$ [µm] | Tn-Trt [min] | (Tn-Trt)/ Td | Tc-Tn [min] | (Tc-Tn)/ Td | Tc-Trt [min] |
|---|---|---|---|---|---|---|---|---|---|
| **STK13** | Acetate | 401 | 660±210 | 1.45±0.2 | 65.6±65.4 | 0.10±0.09 | 39.1±37.5 | 0.06±0.06 | 105±72.2 |
| | Alanine-TrEl | 215 | 213±58.9 | 1.76±0.28 | 25.6±21.7 | 0.12±0.09 | 2.1±10.1 | 0.01±0.05 | 27.7±22.9 |
| | Mannose | 302 | 195±61.5 | 1.51±0.27 | 26±23.9 | 0.13±0.12 | 11.1±12.1 | 0.06±0.06 | 37.1±26.1 |
| | Glycerol | 420 | 165±51 | 1.72±0.23 | 16.2±15.1 | 0.11±0.1 | 12.0±12.2 | 0.07±0.07 | 28.0±18.1 |
| | Glycerol-TrEl | 344 | 148±44 | 1.61±0.18 | 21±21.3 | 0.15±0.16 | 14.6±10.7 | 0.10±0.07 | 35.6±24.2 |
| | Glucose | 259 | 113±31.2 | 1.8±0.23 | 30±18.7 | 0.27±0.16 | 8.8±6.7 | 0.08±0.06 | 38.7±20 |
| | Glycerol-Cas | 250 | 93±32.5 | 1.9±0.34 | 28.5±14.9 | 0.34±0.2 | 8.6±6.7 | 0.1±0.07 | 37.2±15.7 |
| | Glucose-CAS | 409 | 70±23 | 2.01±0.34 | 38±15.7 | 0.58±0.24 | 8.1±9.3 | 0.11±0.11 | 46.1±16.6 |
| | EZRDM | 202 | 37±6 | 3.49±0.51 | 42.5±10.1 | 1.16±0.31 | 0.85±3.7 | 0.02±0.10 | 43.3±10.0 |
| **JM85** | Glycerol | 420 | 159±48 | 2.04±0.3 | NA | NA | NA | NA | 20.6±16.9 |
| | Glucose-Cas | 406 | 74±30 | 2.1±0.4 | NA | NA | NA | NA | 38.3±18.5 |
| **BW27783** (WT) | Glycerol | 108 | 175±50 | 1.72±0.28 | NA | NA | NA | NA | NA |
| **JM111** | Glycerol-TrEl | 300 | 154±44 | 1.43±0.23 | 31.2±33.0 | 0.21±0.21 | 11.8±11.3 | 0.08±0.08 | 43.0±33.0 |
| **JM116** non-polar | Glycerol-TrEl | 318 | 146±52 | 1.92±0.39 | 28.6±27.1 | 0.21±0.20 | 10.6±11.3 | 0.08±0.08 | 39.2±28.2 |
| **JM116** polar | Glycerol-TrEl | 113 | NA | 1.87±0.37 | -6±45.3 | NA | 13.5±16.7 | NA | 7.5±43.8 |
| **JM121** | Glycerol-TrEl | 214 | 156±58 | 1.69±0.25 | 32.9±37.4 | 0.23±0.26 | 15.5±16.0 | 0.10±0.09 | 48.4±40.2 |
| **JM122** | Glycerol-TrEl | 202 | 143±56 | 1.98±0.40 | 35.2±41 | 0.27±0.33 | 14.0±18.4 | 0.11±0.11 | 49.4±37.7 |
| **JM123** | Glycerol-TrEl | 223 | 149±50 | 1.67±0.20 | 29.4±24.9 | 0.20±0.15 | 15.4±13.7 | 0.11±0.08 | 44.8±26.3 |
| **JM127** | Glycerol-TrEl | 251 | 162±63 | 1.74±0.29 | 26.8±30.5 | 0.19±0.24 | 23.3±17.7 | 0.15±0.10 | 50.2±29.5 |

**Table S3.** List of growth media, carbon sources and supplements.

| Media | Buffer | Carbon source, final concentration | Supplements (final concentration) |
|---|---|---|---|
| EZRDM | MOPS | Glucose, 0.5% | 1x ACGU, 1x EZ supplement, $K_2HPO_4$ (1.32 mM) |
| Glucose-Cas | M9 minimal | Glucose, 0.5% | Casamino acids (Cas), 0.2% |
| Glycerol-Cas | M9 minimal | Glycerol, 0.3% | Cas, 0.2% |
| Glucose | M9 minimal | Glucose, 0.5% | |
| Glycerol-TrEl | M9 minimal | Glycerol, 0.3% | 1x Trace metal Elements mix (TrEl) |
| Glycerol | M9 minimal | Glycerol, 0.3% | |
| Mannose | M9 minimal | Mannose, 0.5% | |
| Alanine-TrEl | M9 minimal | Alanine, 0.3% | 1x TrEl |
| Acetate | M9 minimal | Sodium acetate, 0.4% | |

**Table S4.** Correlations involving $Tri$, $Trt$, $Tn$, $C$ and $Td$ in all growth conditions for STK13 strain. Note that the two fastest growth conditions are excluded as the determination of initiation timings in these conditions is ambiguous. Numbers in the parenthesis indicate 95% confidence intervals. For the doubling times and C-periods mean±std is shown.

| **Media** | **No. of cells** | $\boldsymbol{Td}$ **[min]** | $\boldsymbol{C}$ **[min]** | $\boldsymbol{R(C, Td)}$ | **Slope of** $\boldsymbol{Trt}$ **vs** $\boldsymbol{Tri}$ | $\boldsymbol{R(C, Tn - Trt)}$ |
|---|---|---|---|---|---|---|
| Acetate | 401 | 660±210 | 197±64 | 0.63 (0.56, 0.68) | 1.07 (1.04, 1.10) | 0.36 (0.27, 0.44) |
| Alanine-TrEl | 215 | 213±58.9 | 95±28 | 0.65 (0.56, 0.72) | 1.10 (1.02, 1.18) | 0.00 (-0.13, 0.13) |
| Mannose | 298 | 196±61.2 | 101±32 | 0.65 (0.58, 0.71) | 0.91 (0.84, 0.98) | 0.24 (0.13, 0.34) |
| Glycerol | 419 | 165±51 | 88±25 | 0.70 (0.65, 0.75) | 1.08 (1.02, 1.14) | -0.07 (-0.16, 0.03) |
| Glycerol-TrEl | 344 | 148±44 | 85±18 | 0.64 (0.57, 0.70) | 1.10 (1.05, 1.14) | -0.15 (-0.25, -0.05) |
| Glucose | 259 | 113±31.2 | 76±15 | 0.67 (0.59, 0.73) | 0.98 (0.90, 1.06) | -0.02 (-0.15, 0.10) |
| Glycerol-Cas | 250 | 93±32.5 | 63±15 | 0.70 (0.63, 0.76) | 0.96 (0.87, 1.05) | -0.17 (-0.29, -0.05) |

**Table S5.** Comparing delay time between termination of replication and initiation of constriction ($Tn - Trt$) in WT and mutant cells. The table lists $p$-values from Mann-Whitney and t-test.

| **Tn-Trt WT vs mutant strains** | **JM116 (MinC) NonPolar** | **JM116 (MinC) Polar** | **JM121 (SlmA)** | **JM123 (MatP)** | **JM122 (ZapA)** | **JM127 (ZapB)** | **JM111 (FtsK K997A)** |
|---|---|---|---|---|---|---|---|
| Mann-Whitney | 2.13E-05 | 2.56E-22 | 3.58E-04 | 4.63E-06 | 1.01E-04 | 0.0713 | 1.24E-04 |
| t-test | 5.17E-05 | 2.83E-16 | 2.00E-06 | 1.77E-05 | 1.29E-07 | 0.0054 | 2.47E-06 |
